# RPE pseudopods maintain photoreceptor outer segment renewal despite subretinal space expansion in *Adam9* knockout mice

**DOI:** 10.64898/2025.12.16.694731

**Authors:** Tylor R. Lewis, Carson M. Castillo, Sebastien Phan, Camilla R. Shores, Kylie K. Hayase, Keun-Young Kim, Mark H. Ellisman, Oleg Alekseev, Marie E. Burns, Vadim Y. Arshavsky

## Abstract

Vision begins in the outer segment compartment of photoreceptor cells, which is constantly renewed through the addition of membrane material at its base and ingestion of mature membranes at its tip by the retinal pigment epithelium (RPE). The close apposition of outer segments to the RPE is believed to be critical for maintaining this renewal process. Yet, in several retinal diseases, expansion of the subretinal space separating photoreceptors from the RPE does not immediately impact photoreceptor functionality. Here, we analyzed outer segment function and renewal in the *Adam9* knockout mouse characterized by a major expansion of the subretinal space. Surprisingly, photoreceptor-RPE separation affected neither the sensitivity of photoreceptor light-responses nor the normal rate of outer segment renewal in this mouse prior to the onset of photoreceptor degeneration. The latter is achieved through the formation of elongated RPE “pseudopods” extending across the enlarged subretinal space to ingest outer segment tips. This work suggests that pseudopod formation may underlie the persistence of photoreceptor function in human diseases accompanied by photoreceptor-RPE separation, such as vitelliform macular dystrophy or age-related macular degeneration associated with subretinal drusenoid deposits.

## Introduction

Vertebrate vision begins with absorption of light by the specialized ciliary outer segment compartment of photoreceptor cells. The outer segment contains a stack of hundreds to thousands of light-sensitive disc-shaped membranes, which is continuously renewed by addition of new discs at the base and ingestion of old discs at the tip by the apposing retinal pigment epithelium (RPE) (1). This process is critical for maintaining photoreceptor health, as mutations in proteins involved in outer segment renewal are associated with multiple retinal degenerative diseases (2, 3).

Outer segment turnover does not occur in retinas detached from the RPE (4) and it is widely believed that the close apposition between outer segments and the RPE is essential for maintaining photoreceptor health (e.g. (3)). Yet in diseases causing various degrees of subretinal space expansion, such as vitelliform macular dystrophy or age-related macular degeneration associated with subretinal drusenoid deposits, the defects in visual function and photoreceptor morphology are much less pronounced than one would expect from the observed degree of separation between the retina and RPE (e.g. (5, 6)). The minimal impact of subfoveal deposits on patients’ best-corrected visual acuity is a well-recognized clinical feature of Best vitelliform macular dystrophy (VMD2 aka Best disease; Online Mendelian Inheritance in Man #153700) in its vitelliform stage that can last for years prior to the subsequent degenerative stage (e.g. (7)). These observations in human patients suggest that outer segment turnover may be maintained in the absence of a close apposition between outer segments and the RPE. We tested this hypothesis by analyzing the dynamics of outer segment renewal in *Adam9* knockout (*Adam9^−/−^*) mice.

Mutations in the *ADAM9* gene cause retinal degenerative disease in humans and dogs (8–10). Interestingly, a separation between outer segments and the RPE associated with a disruption in the structure of RPE apical microvilli has been observed in both affected dogs (9) and a mouse model of ADAM9-associated disease (8). It was also found that the subretinal space in *Adam9^−/−^*mice contains mononuclear phagocytes (8). This raises an intriguing possibility that the role of the RPE in ingesting outer segment tips in these animals is instead performed by these phagocytic cells.

We now show that the gap between outer segments and the RPE in *Adam9^−/−^*mice forms early in development, significantly prior to the onset of photoreceptor degeneration. Despite the disruption of this cellular interface, photoreceptors function normally and maintain a normal rate of outer segment renewal prior to the onset of their degeneration. The latter is achieved by RPE cells forming elongated “pseudopods” that reach across the expanded subretinal space of *Adam9^−/−^* mice to contact and ingest outer segment tips. Notably, the mononuclear phagocytes that massively migrate to the subretinal space do not appear to contribute to this process. The depletion of these cells from the *Adam9^−/−^*retina did not have any appreciable effect on either photoreceptor function or survival. Overall, these data reveal that normal photoreceptor outer segment renewal can occur in the absence of a close apposition between photoreceptors and the RPE. Furthermore, these findings explain how the RPE could support outer segment turnover in human diseases associated with an expansion of the subretinal space.

## Results

### Photoreceptor function in human vitelliform macular dystrophy is minimally affected

To highlight the impact of a pathological expansion of the subretinal space on photoreceptor function in humans, we present a multimodal assessment of a patient afflicted with vitelliform macular dystrophy type 2 caused by a pathogenic variant in *BEST1* (c.17C>G, p.Thr6Arg) (11–13) (Figure 1). This asymptomatic 12-year-old female was evaluated due to a positive family history of macular disease. Imaging with optical coherence tomography (Figure 1A) and fundus autofluorescence photography (Figure 1B) revealed the presence of bilateral central vitelliform lesions. Despite the significant separation of subfoveal photoreceptors from the RPE, the patient’s best-corrected visual acuity was logMAR 0.2 (Snellen equivalent 20/32) in each eye. Her central cone sensitivity was only minimally reduced on microperimetry testing (Figure 1C) and foveolar cone function was only borderline attenuated on multifocal electroretinography (zone 1 in Figure 1D). These observations, consistent with other reports in patients having expansions in the subretinal space (e.g. (5–7)), led us to evaluate the degree to which a close apposition between photoreceptor outer segments and the RPE is essential for supporting photoreceptor cell function.

**Figure 1.**
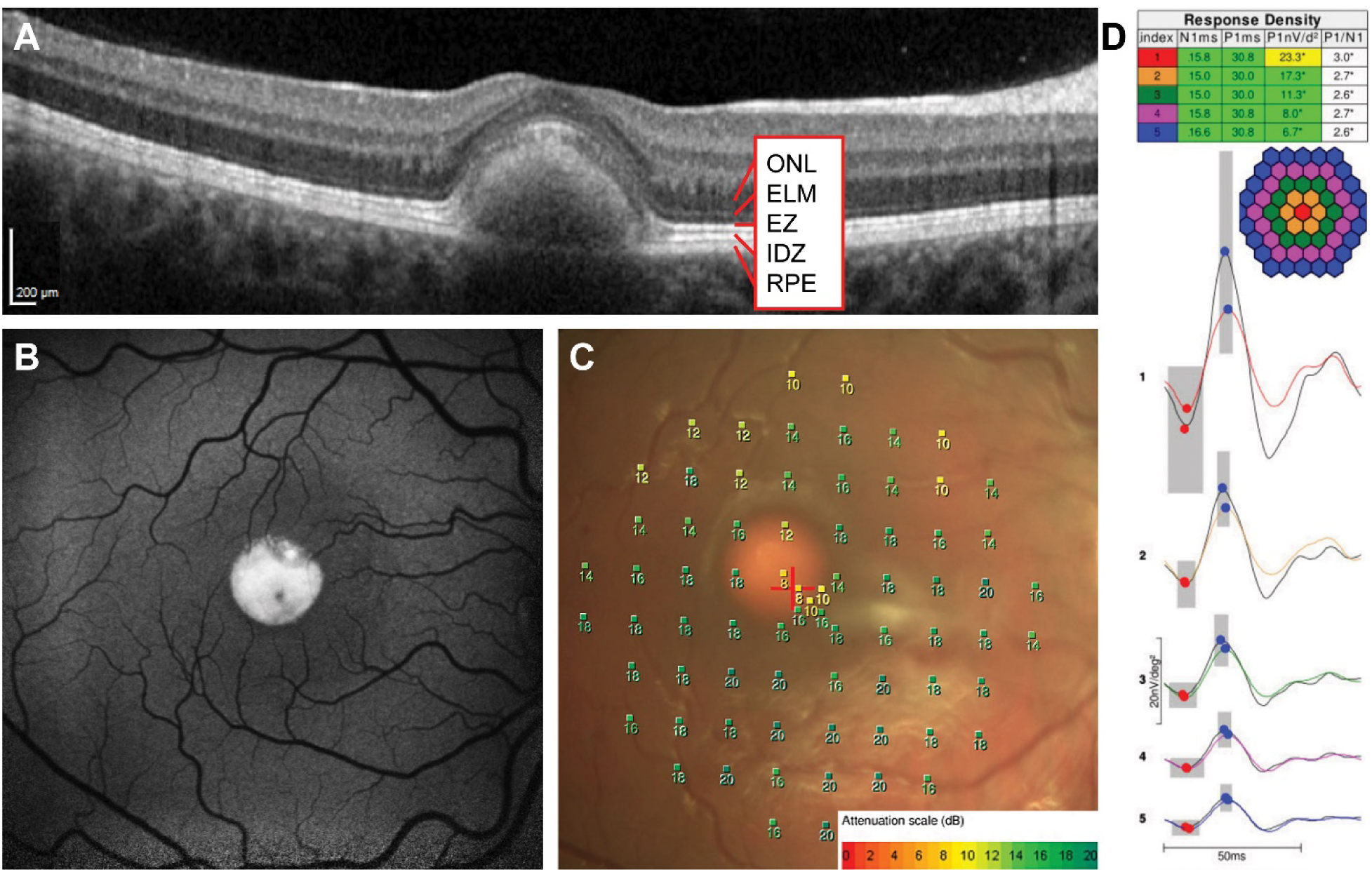
Vitelliform macular dystrophy is characterized by a separation of photoreceptors and the RPE without major early visual defects. Right eye of a 12-year-old patient with a subfoveal vitelliform lesion due to vitelliform macular dystrophy type 2. (A) Spectral-domain optical coherence tomography image through the foveola. ONL: outer nuclear layer; ELM: external limiting membrane; EZ: ellipsoid zone; IDZ: interdigitation zone; RPE: retinal pigment epithelium. (B) Fundus autofluorescence photograph of the macula. (C) Microperimetry measurements at 73 macular locations superimposed on the corresponding fundus color photograph. The sensitivity scale (in dB) is shown at the bottom. (D) Multifocal electroretinography recordings obtained to a stimulus array of 61 elements. Individual responses were averaged within five different eccentricity ranges (color-coded in the hexagon schematic) and arranged vertically from center to periphery. Averaged response density data are tabulated above. Grey boxes represent normal ranges of response amplitude and timing (vertical and horizontal dimensions, respectively).

### Photoreceptor loss in the ADAM9 knockout mouse does not occur until after P60

To address the requirement for a close apposition between photoreceptors and the RPE in maintaining photoreceptor cells, we analyzed the *Adam9^−/−^* mouse (14) that had previously been established to display a separation between outer segments and the RPE at advanced stages of photoreceptor degeneration (8). Our first goal was to assess the time course of photoreceptor degeneration occurring in this mouse model, as the previous study (8) only analyzed *Adam9^−/−^* mice of 1 year of age or older. We analyzed the retinal morphology of younger *Adam9^−/−^* mice and found no signs of photoreceptor degeneration at either P30 or P60 (Figure 2). However, retinal degeneration in *Adam9^−/−^* mice took place at later ages, as ∼30% of photoreceptors were lost by P180 and ∼50% were lost by P365.

**Figure 2.**
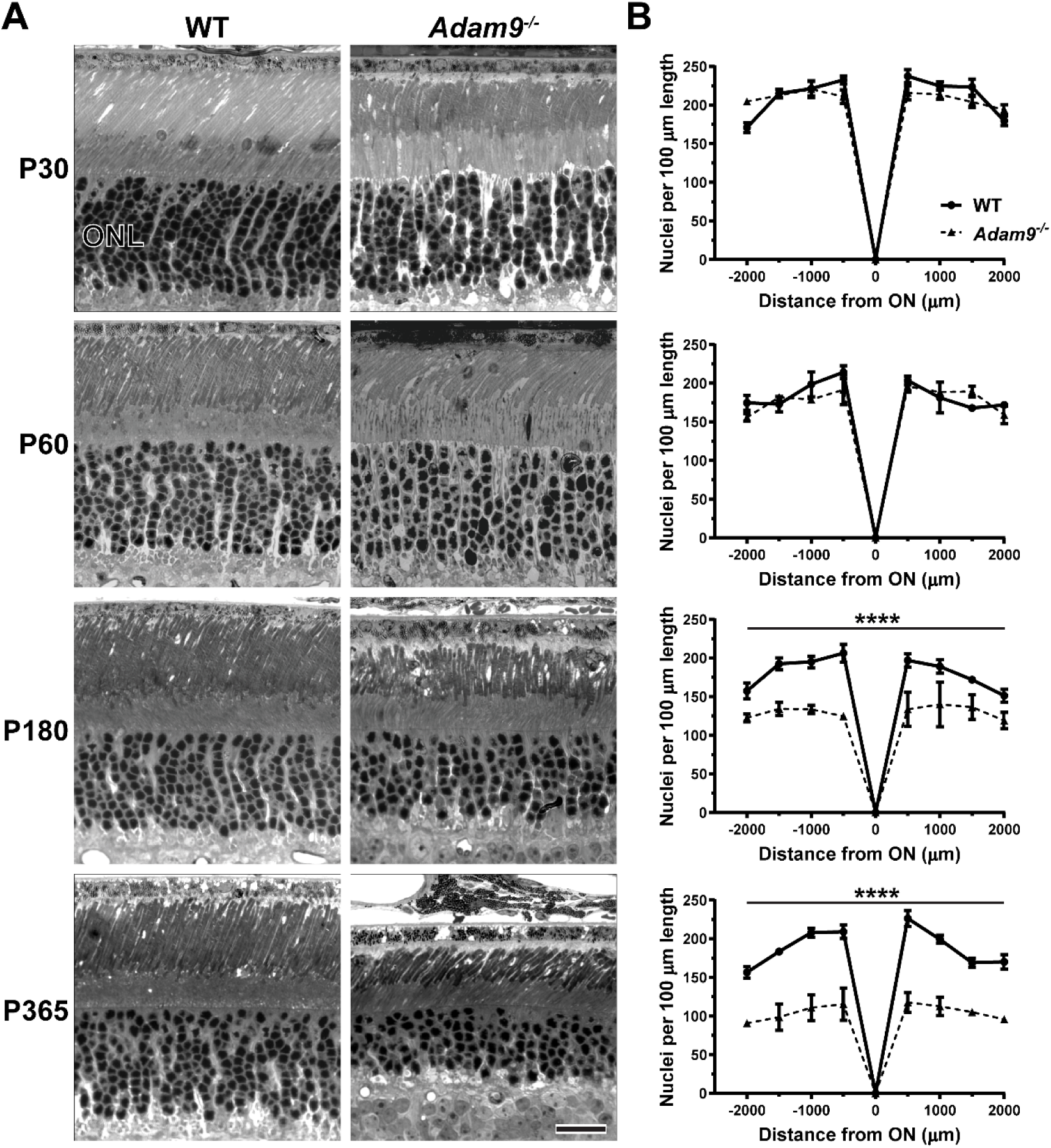
Loss of ADAM9 leads to a slow, progressive photoreceptor degeneration. **(A)** Representative light microscopy images of WT and *Adam9^−/−^* retinas at P30 through P365. ONL: outer nuclear layer. Scale bar: 20 µm. **(B)** Quantification of the number of photoreceptor nuclei in the ONL in 100 µm segments of retinal sections located at 500 µm intervals from the optic nerve (ON) at each timepoint. For each genotype, three retinas from separate mice were analyzed. Two-way ANOVA was used to compare the nuclear counts (not including optic nerve) between genotypes at each timepoint. At P30 and P60, there was no statistically significant reduction in photoreceptor numbers in *Adam9^−/−^* retinas compared to WT retinas (p = 0.3443 and 0.2870, respectively). At each subsequent timepoint, there was a statistically significant reduction in photoreceptor numbers in *Adam9^−/−^* retinas compared to WT retinas (p < 0.0001 for both P180 and P365). Error bars represent mean ± SEM.

We next analyzed photoreceptor function using electroretinography (ERG) (Figure 3). While there were no overt defects in ERG responses of *Adam9^−/−^* mice at either P30 or P60, there was a decrease in the amplitudes of ERG a-wave responses in older *Adam9^−/−^* mice, with an ∼30% reduction observed at P180 and ∼50% reduction at P365. Because the ERG a-waves represent the compounded light responses of photoreceptor cells, a nearly perfect correspondence between the reduction in a-wave amplitudes and the number of surviving photoreceptors argues that the surviving photoreceptors remain fully functional at these ages. ERG b-wave responses, reflecting the activity of downstream bipolar cells, were affected as well, although to a lesser degree than a-waves. This is consistent with the underlying pathology originating from photoreceptors.

**Figure 3.**
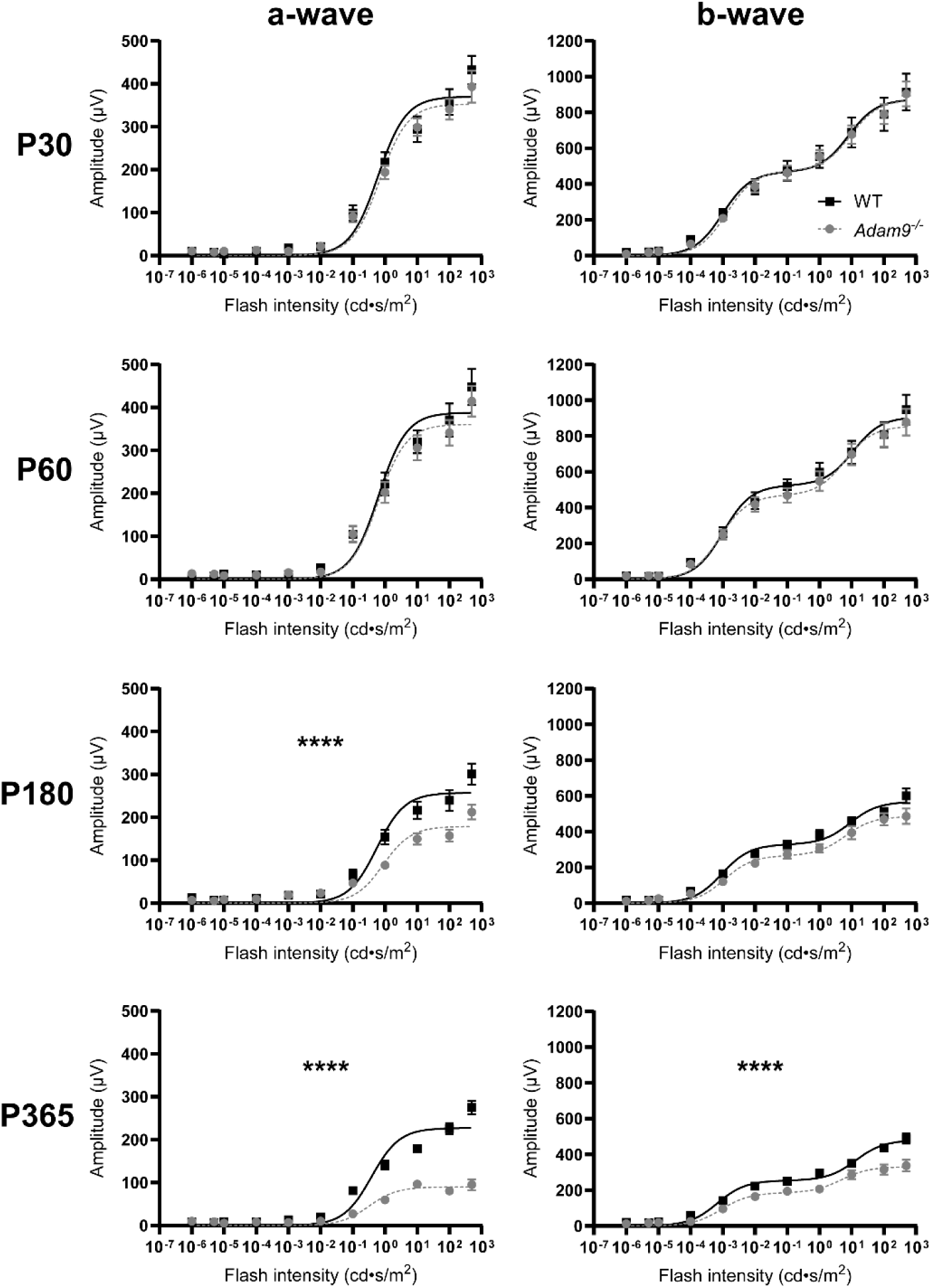
Loss of ADAM9 leads to a slow, progressive loss of visual function. ERG responses of dark-adapted WT and *Adam9^−/−^* mice at P30 through P365 are plotted as a function of flash intensity and fit using a double- or single-hyperbolic function. Scotopic a-wave amplitudes are shown on the left and scotopic b-wave amplitudes are shown on the right. For each genotype at each timepoint, at least 12 eyes were analyzed. Two-way ANOVA was used to determine whether there were any statistically significant differences in genotype/light intensity. At P30 and P60, there were no statistically significant differences in a-waves (p = 0.9622 and 0.9833, respectively) or b-waves (p > 0.9999 and 0.9987, respectively). At P180, there was a statistically significant difference in a-wave (p < 0.0001) but not b-wave (p = 0.9987). At P365, there was a statistically significant difference in both a-wave (p < 0.0001) and b-wave (p < 0.0001). Error bars represent mean ± SEM.

Further evidence that photoreceptors of *Adam9^−/−^* mice remain functionally intact before they degenerate was obtained in single cell recordings from their rods at P60 (Table 1). The average dark currents (I_dark_) and flash sensitivities (I_o_) were statistically indistinguishable from those obtained in photoreceptors of WT mice. The dim flash response kinetics of *Adam9^−/−^* rods, as measured by the time to peak, time constant of recovery (_Trec_) and integration time, were also not different from those of WT rods. These findings indicate that rods of *Adam9^−/−^* mice display a normal ability to produce responses to light.

**Table 1.** Single cell suction electrode recordings of WT and *Adam9^−/−^* rods. Values are mean ± SEM (n = number of cells) for each of the single cell response parameters, as defined in the Methods.

| | Dark<br>current<br>$I_{\text{dark}}$<br>(pA) | Flash<br>Sensitivity<br>$I_0$<br>(photons/$\mu\text{m}^2$) | Dim<br>flash<br>Time to<br>peak<br>(ms) | Dim flash<br>recovery<br>$\tau_{\text{rec}}$<br>(ms) | Dim flash<br>Integration<br>time (ms) | Saturating<br>flash<br>recovery<br>$\tau_D$ (ms) |
| --- | --- | --- | --- | --- | --- | --- |
| **WT** | 13.5 ± 0.7<br>(21) | 54 ± 4<br>(21) | 103 ± 8<br>(21) | 152 ± 10<br>(19) | 239 ± 20<br>(21) | 190 ± 15<br>(19) |
| *Adam9*<sup>-/-</sup> | 12.0 ± 1.9<br>(6) | 44 ± 6<br>(5) | 125 ± 15<br>(6) | 161 ± 23<br>(5) | 254 ± 29<br>(6) | 164 ± 13<br>(5) |

### Subretinal space expansion in the ADAM9 knockout mouse occurs early in photoreceptor development

We next used transmission electron microscopy (TEM) to analyze the ultrastructure of the outer segment-RPE interface of *Adam9^−/−^* mice (Figure 4). Strikingly, we found that the disruption in the normally close apposition between outer segments and the RPE, previously reported for aged mice (8), begins as early as P14. At this age, the subretinal space is expanded and filled with RPE microvilli that do not ensheathe outer segments. As *Adam9^−/−^* mice age, the gap between the RPE and outer segment tips widens and the network of RPE microvilli expands. Many of these microvilli appear to be swollen and some contain pigment granules displaced from the RPE somas. Of note, immunofluorescent imaging of RPE flatmounts from *Adam9^−/−^*mice did not reveal any overt pathology in the shape or structure of the RPE monolayer at P30 or P60, indicating that the expanded subretinal space is not immediately associated with RPE degeneration (Supplemental Figure 1).Importantly, the onset of the outer segment-RPE interface disruption precedes any signs of photoreceptor degeneration by over 6 weeks. This suggests that a close apposition between outer segments and the RPE is not required for maintaining photoreceptor health over a long period of time.

**Figure 4.**
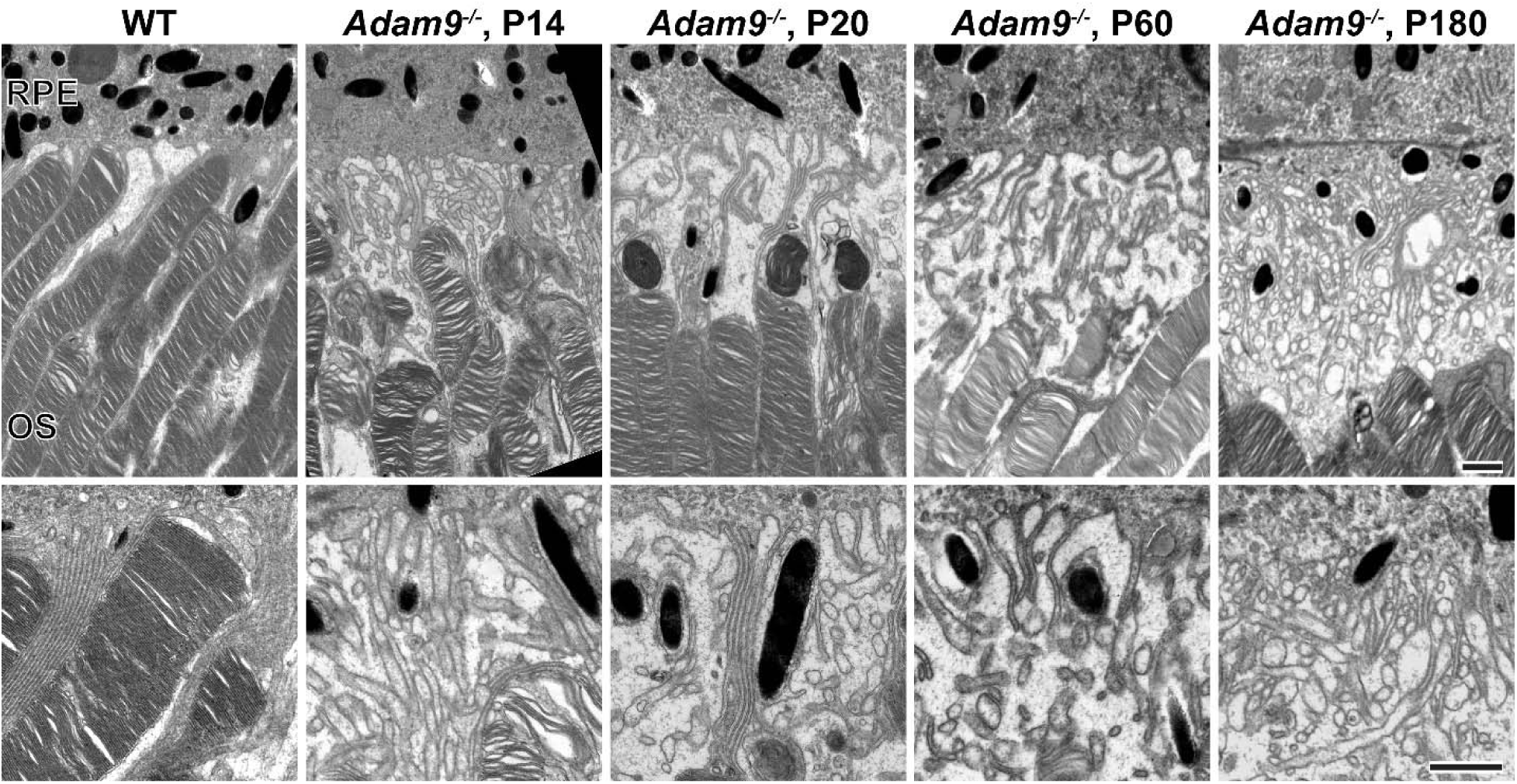
Loss of ADAM9 leads to an early separation of the outer segment-RPE interface. Representative TEM images of the photoreceptor outer segment/RPE interface in WT (P30) and *Adam9^−/−^* (P14 through P180) mice. Higher magnification images are shown in the bottom panels. OS: outer segment. Scale bars: 1 µm.

### Subretinal space expansion in the ADAM9 knockout mouse does not affect normal outer segment renewal

We next sought to determine whether the separation between photoreceptors and the RPE in *Adam9^−/−^* mice affects the process of outer segment renewal. We first analyzed whether outer segment volume was altered in these mice prior to the onset of photoreceptor degeneration. Because rhodopsin represents >90% of the protein material within the outer segment membranes, its retinal content serves as a sensitive measure of the total outer segment volume. We determined the rhodopsin content in the retinas of *Adam9^−/−^* mice before the onset of photoreceptor degeneration using difference spectroscopy and found it to be the same as in WT mice (Supplemental Figure 2). Considering the unchanged total number of rods between these mice and the unchanged total rhodopsin content, we conclude that the volume of individual rod outer segments was unaffected by the loss of ADAM9.

We next analyzed the dynamics of outer segment renewal in *Adam9^−/−^* mice using a recently described approach (15). This approach takes advantage of the fact that the newly forming discs at the rod outer segment base are exposed to the extracellular space before becoming enclosed within the outer segment plasma membrane in the final step of their maturation (reviewed in (16)). Accordingly, these discs can entrap a membrane-impermeable fluorescent dye, CF-568-hydrazide, transiently introduced to the eye via an intravitreal injection. As outer segment renewal continues, discs containing the entrapped dye migrate outward along the outer segment axis. At any given time, the distance between the base of the outer segment and the dye band reflects the amount of new outer segment material added since the time of injection.

We intravitreally injected WT and *Adam9^−/−^* mice with CF-568-hydrazide at P30 when the gap between outer segments and the RPE is already present but before the onset of photoreceptor degeneration. At 6 days post-injection, outer segments were isolated, stained with wheat germ agglutinin (WGA) and imaged (Figure 5). In both groups of P30 mice, a small band of the CF-568-hydrazide dye was observed at ∼60% of the outer segment length from the base, reflecting the rate of outer segment renewal in WT mouse rods (∼10% per day) established by multiple approaches (1), including previous experiments with CF-568-hydrazide in adult mice (17). Of note, following outer segment isolation for imaging, there appeared to be a small ∼15% decrease in their overall length in *Adam9^−/−^* mice. Considering the essentially normal volume of outer segments (as assumed from the total rhodopsin content above), this likely suggests that *Adam9^−/−^* outer segments may be more susceptible to mechanical damage at their tips. The normal rate of outer segment morphogenesis in P30 *Adam9^−/−^* mice suggests that the removal of old discs at the outer segment tips persists essentially normally despite the ongoing expansion of the subretinal space.

**Figure 5.**
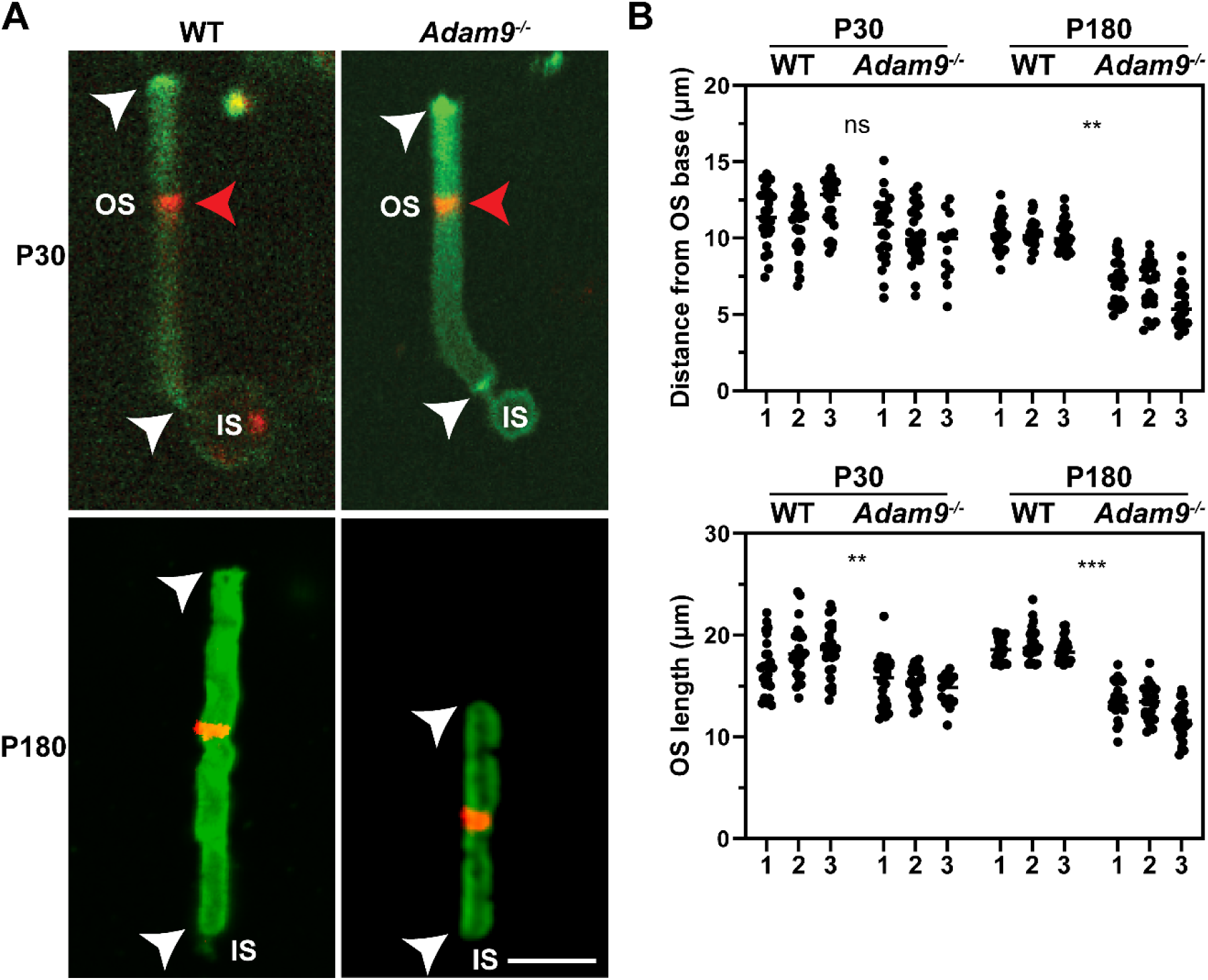
Loss of ADAM9 does not affect the dynamics of outer segment renewal before the onset of photoreceptor degeneration. **(A)** Photoreceptor cells from WT and *Adam9^−/−^* mice isolated 6 days after intravitreal injection of CF-568-hydrazide (red) and stained with WGA (green). The distance between the red band in the outer segment (OS) and its base near the inner segment (IS) reflects the new outer segment material formed within the time period following the injection. White arrowheads indicate the ends of the OS; red arrowheads indicate the CF-568-hydrazide band. Scale bar: 5 µm. **(B)** Quantification of the position of the CF-568-hydrazide band as the distance from the outer segment base and quantification of the outer segment length. Individual points represent isolated cells from one of three different injected eyes (indicated at the bottom) for each genotype and age. The number of cells analyzed for each experiment were: WT, P30 1: 25; WT, P30 2: 25; WT, P30 3: 25; *Adam9^−/−^*, P30 1: 25; *Adam9^−/−^*, P30 2: 25; *Adam9^−/−^*, P30 3: 13; WT, P180 1: 25; WT, P180 2: 25; WT, P180 3: 25; *Adam9^−/−^*, P180 1: 25; *Adam9^−/−^*, P180 2: 23; *Adam9^−/−^*, P180 3: 22. Unpaired t-test, using the average value from each of the three eyes, showed no statistically significant difference between the location of the dye band in WT and *Adam9^−/−^* mice at P30 (p = 0.0678), but a statistically significant difference at P180 (p = 0.0018). For outer segment length, there was a statistically significant difference between WT and *Adam9^−/−^* mice at P30 (p = 0.0073) and at P180 (p = 0.0009). Line represents mean.

We next used this approach to analyze outer segment renewal at P180, following the onset of photoreceptor degeneration in *Adam9^−/−^* mice (Figure 5). Strikingly, we found that, in this case, outer segment renewal of P180 *Adam9^−/−^* mice was slowed by ∼40% and is accompanied by an ∼30% reduction in outer segment length.

### RPE cells form extensive pseudopods to ingest outer segment tips in the ADAM9 knockout mouse

Our finding that, prior to photoreceptor degeneration, the rate of outer segment renewal is unaffected by the ADAM9 knockout despite the subretinal space expansion led us to explore the mechanism supporting the unaltered rate of outer segment turnover. We envisioned two possibilities: (1) the RPE retains the ability to ingest outer segment material despite a separation between these two cell types, or (2) mononuclear phagocytes infiltrating the subretinal space (8) may instead fulfill this role. We first tested the role of the RPE by quantifying the number of outer segment-containing phagosomes in the RPE of WT and *Adam9^−/−^* mice at the diurnal peak of phagosome accumulation, which occurs shortly after light onset. We found a small, yet not statistically significant reduction in phagosome number and no change in phagosome size (Figure 6, A and B). This suggests that RPE cells retain a nearly normal ability to ingest outer segment material, despite the lack of a close apposition to photoreceptor cells. Intriguingly, phagosome number was significantly decreased at P180 after the onset of photoreceptor degeneration, which is consistent with our observation of decreased outer segment renewal at this timepoint.

**Figure 6.**
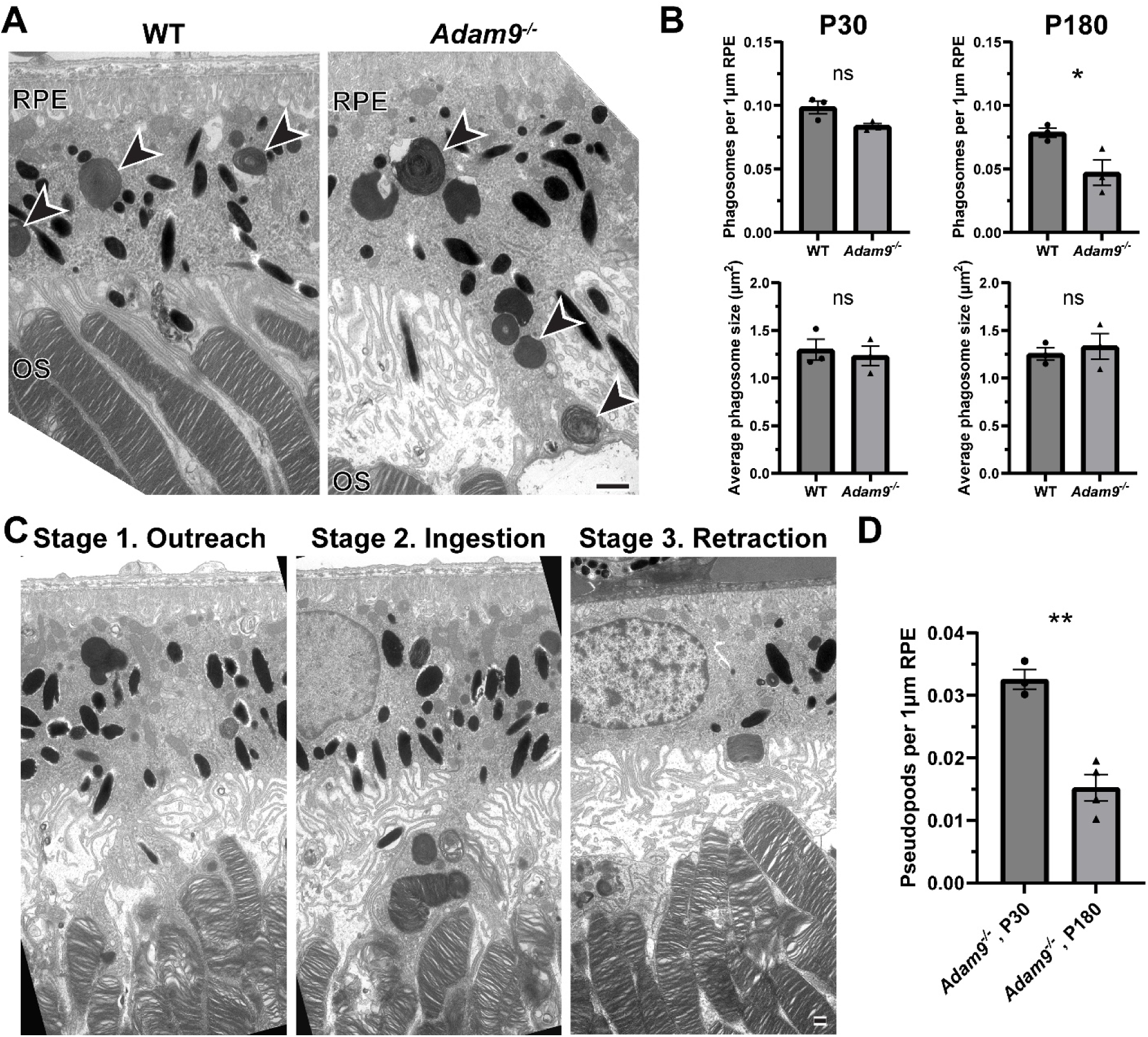
Loss of ADAM9 leads to the formation of elongated RPE pseudopods supporting outer segment renewal. **(A)** Representative TEM images of the outer segment/RPE interface in P30 WT and *Adam9^−/−^*mice. Black arrowheads indicate phagosomes. Scale bar: 1 µm. **(B)** Quantification of phagosome number (normalized to RPE length) and phagosome size at P30 and P180. For each condition, three eyes were analyzed. For number of phagosomes, the RPE length analyzed was: WT, P30 1: 497 µm; WT, P30 2: 520 µm; WT, P30 3: 420 µm; *Adam9^−/−^*, P30 1: 445 µm; *Adam9^−/−^*, P30 2: 433 µm; *Adam9^−/−^*, P30 3: 543 µm; WT, P180 1: 566 µm; WT, P180 2: 646 µm; WT, P180 3: 652 µm; *Adam9^−/−^*, P180 1: 642 µm; *Adam9^−/−^*, P180 2: 604 µm; *Adam9^−/−^*, P180 3: 629 µm. Unpaired t-test showed a statistically significant difference in phagosome number between WT and *Adam9^−/−^* mice at P180 (p = 0.0430) but not P30 (p = 0.0543). For phagosome size, 25 phagosomes from each sample were analyzed. Unpaired t-test showed no statistically significant difference between phagosome size of WT and *Adam9^−/−^*mice at either P30 (p = 0.6883) or P180 (p = 0.6173). **(C)** Representative TEM images of pseudopods in P30 *Adam9^−/−^* mice at various stages of their formation. Scale bar: 1 µm. **(D)** Quantification of the pseudopod number normalized to the RPE length analyzed. For each timepoint, at least three eyes were analyzed. The RPE length analyzed was: *Adam9^−/−^*, P30 1: 437 µm; *Adam9^−/−^*, P30 2: 394 µm; *Adam9^−/−^*, P30 3: 464 µm; *Adam9^−/−^*, P180 1: 390 µm; *Adam9^−/−^*, P180 2: 409 µm; *Adam9^−/−^*, P180 3: 394 µm, *Adam9^−/−^*, P180 4: 447 µm. Unpaired t-test showed a statistically significant reduction in pseudopod number at P180 (p = 0.0016). Error bars represent mean ± SEM.

Strikingly, *Adam9^−/−^* mice displayed elongated “pseudopods” extending from the RPE cell bodies to contact and ingest outer segment tips (Figure 6A). These pseudopods appeared to be dynamic structures captured in three different stages illustrated in Figure 6C. Some of them extended down to contact outer segment tips without containing phagosomes (Stage 1. “Outreach”), others were enriched with phagosomes (Stage 2. “Ingestion”) and some of them appeared in a state of retraction in which they contained phagosomes but no longer contacted outer segment tips (Stage 3. “Retraction”). In this analysis, we observed a pseudopod approximately every 30 µm of RPE length at P30 and about half as many at P180 (Figure 6D). The structure of these pseudopods can be particularly well-appreciated in three-dimensional electron tomograms spanning ∼3 *μ*m of retinal depth (Movies 1 and 2; Figure 7).

**Figure 7.**
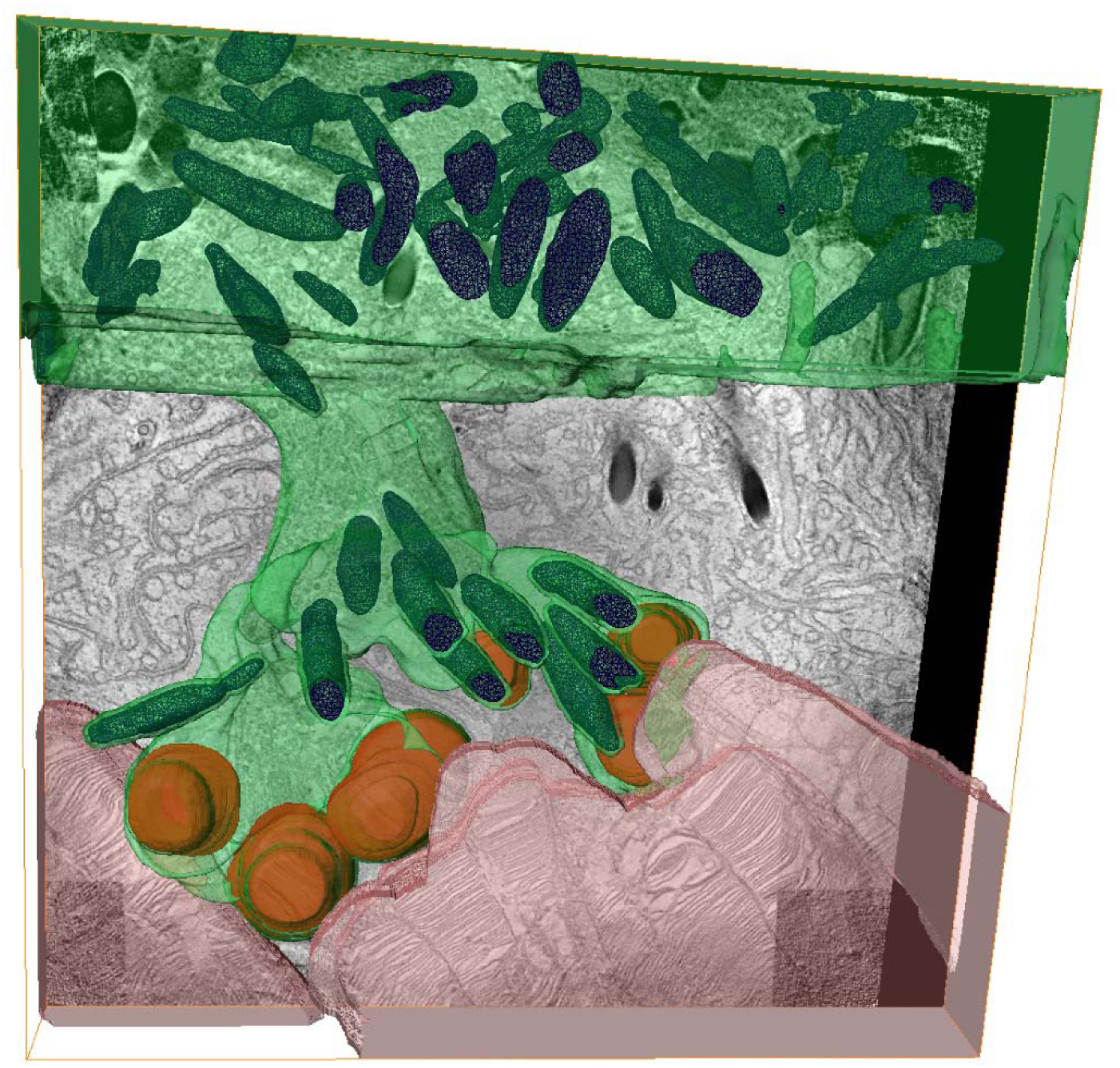
Three-dimensional reconstruction of an RPE pseudopod reaching across the expanded subretinal space of an ADAM9 knockout mouse. Segmentation of three-dimensional electron tomograms (shown in Movie 1) obtained from four 500 nm thick serial sections across the retina of an *Adam9^−/−^* mouse. The RPE is pseudo-colored in green with pigment granules in blue. Outer segments are pseudo-colored in pink. Phagosomes containing outer segment material are pseudo-colored in brown. For clarity, the RPE microvilli are omitted from the segmentation. One individual *z*-section from the tomogram is shown in the background.

RPE pseudopods, also called “phagocytic cups”, have been previously observed in the process of outer segment ingestion in normal retinas (e.g. (18, 19)). Our current data demonstrate that the RPE has an innate ability to form much larger pseudopods to ingest outer segment tips across a pathologically expanded subretinal space. The formation of these pseudopods underlies the normal outer segment renewal in young *Adam9* knockout mice, thereby prolonging the survival of photoreceptor cells. The subsequent slowdown in outer segment renewal and the onset of photoreceptor degeneration in older *Adam9* knockout mice is associated with a reduction in pseudopod density.

### Activated subretinal mononuclear phagocytes do not appear to modulate the pathology associated with loss of ADAM9

In another set of experiments, we explored whether the mononuclear phagocytes accumulating in the subretinal space of *Adam9^−/−^* mice (Figure 8A) may contribute to supporting outer segment renewal and maintaining photoreceptor health. Subretinal immune cells have been observed in many models of retinal degeneration, and it has been debated whether they serve a protective, harmful or perhaps even neutral role (reviewed in (20)). We found that in the *Adam9^−/−^* retinas, the IBA1-positive mononuclear phagocytes are also CD68-positive (Figure 8B), suggesting that they are activated (21). These cells are observed in the subretinal space as early as P30 and densely populate this area by P60 (Figure 8, C and D). Interestingly, their migration to this space occurs at least one month prior to any observable photoreceptor degeneration. Once in the subretinal space, these phagocytes are closely interdigitated with the photoreceptor outer segments and contain ingested outer segment material (Figure 8A and Supplemental Figure 3). Since these immune cells have the capacity to ingest outer segment material, they could, in principle, contribute to the process of outer segment renewal.

**Figure 8.**
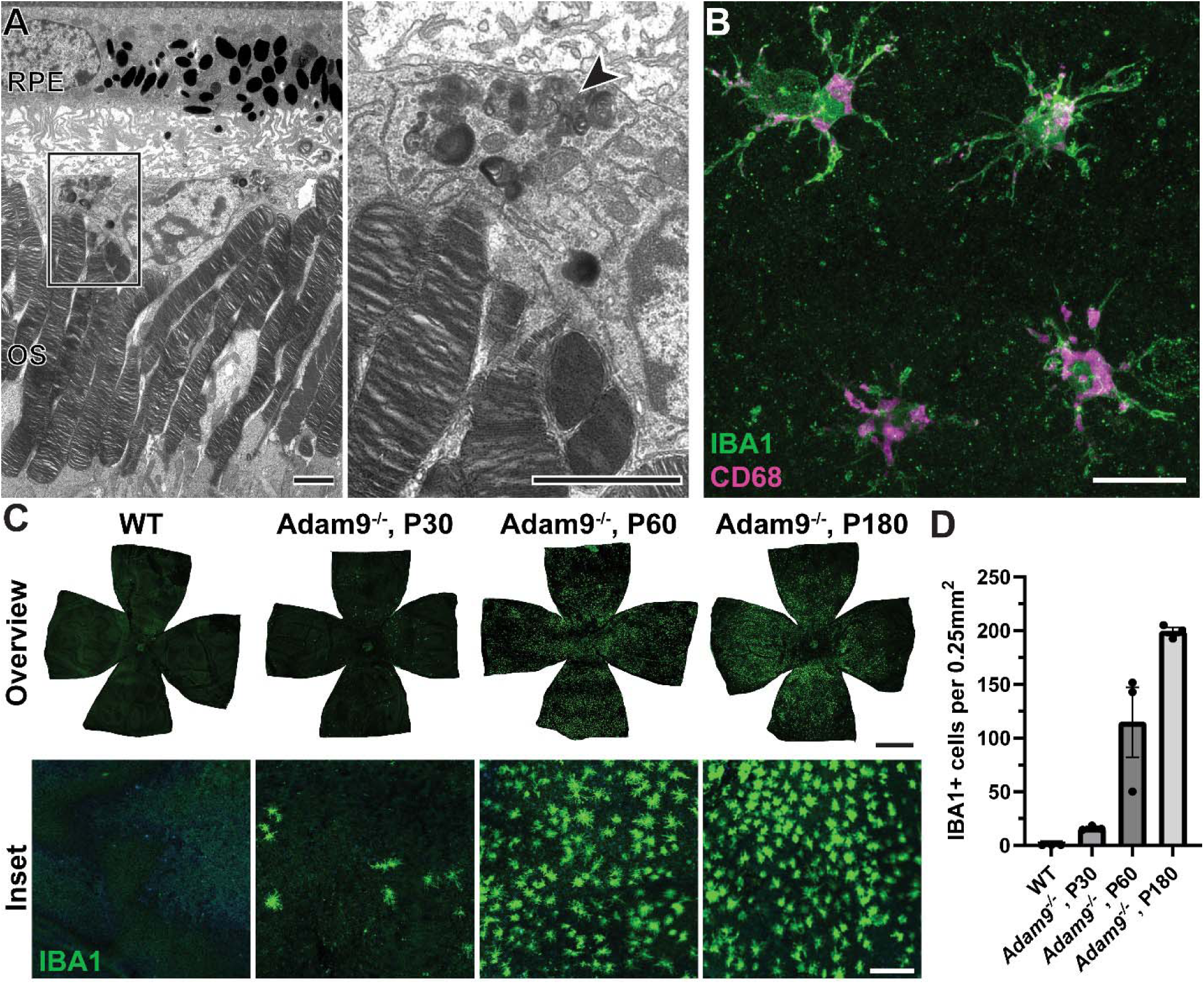
Activated mononuclear phagocytes infiltrate the subretinal space of ADAM9 knockout mice prior to photoreceptor degeneration. **(A)** Representative TEM image of a mononuclear phagocyte in the subretinal space of an *Adam9^−/−^*mouse at P30. Boxed inset is shown on the right. Arrowhead indicates ingested outer segment material inside. OS: outer segment. Scale bars: 2.5 µm. **(B)** Representative immunofluorescence images of RPE flatmounts of an *Adam9^−/−^*mouse at P60 stained with IBA1 (green), a marker for mononuclear phagocytes, and CD68 (magenta), a marker highly enriched in activated microglia in the retina (33). Scale bar: 20 µm. **(C)** RPE flatmounts of WT (P30) and *Adam9^−/−^*(P30-P180) mice stained with IBA1. Whole flatmounts are shown in the overview on top, high magnification insets are shown on bottom. Scale bars: 1,000 µm (top), 100 µm (bottom). **(D)** Quantification of the number of IBA+ cells in a 500 µm x 500 µm area averaged across four 500 µm intervals from the optic nerve (dorsal, nasal, ventral, temporal). For each group, three eyes were analyzed.

To assess the contribution of mononuclear phagocytes in modulating the pathology in *Adam9^−/−^* mice, we depleted them using PLX3397, a CSF1R inhibitor that nearly completely eliminates all myeloid cells from mouse nervous tissues (22), including the retina (23). We first verified the efficacy of this treatment by showing that PLX3397 administered to *Adam9^−/−^* mice starting at P60 depleted the subretinal IBA1-positive cells within 30 days (Figure 9A).

**Figure 9.**
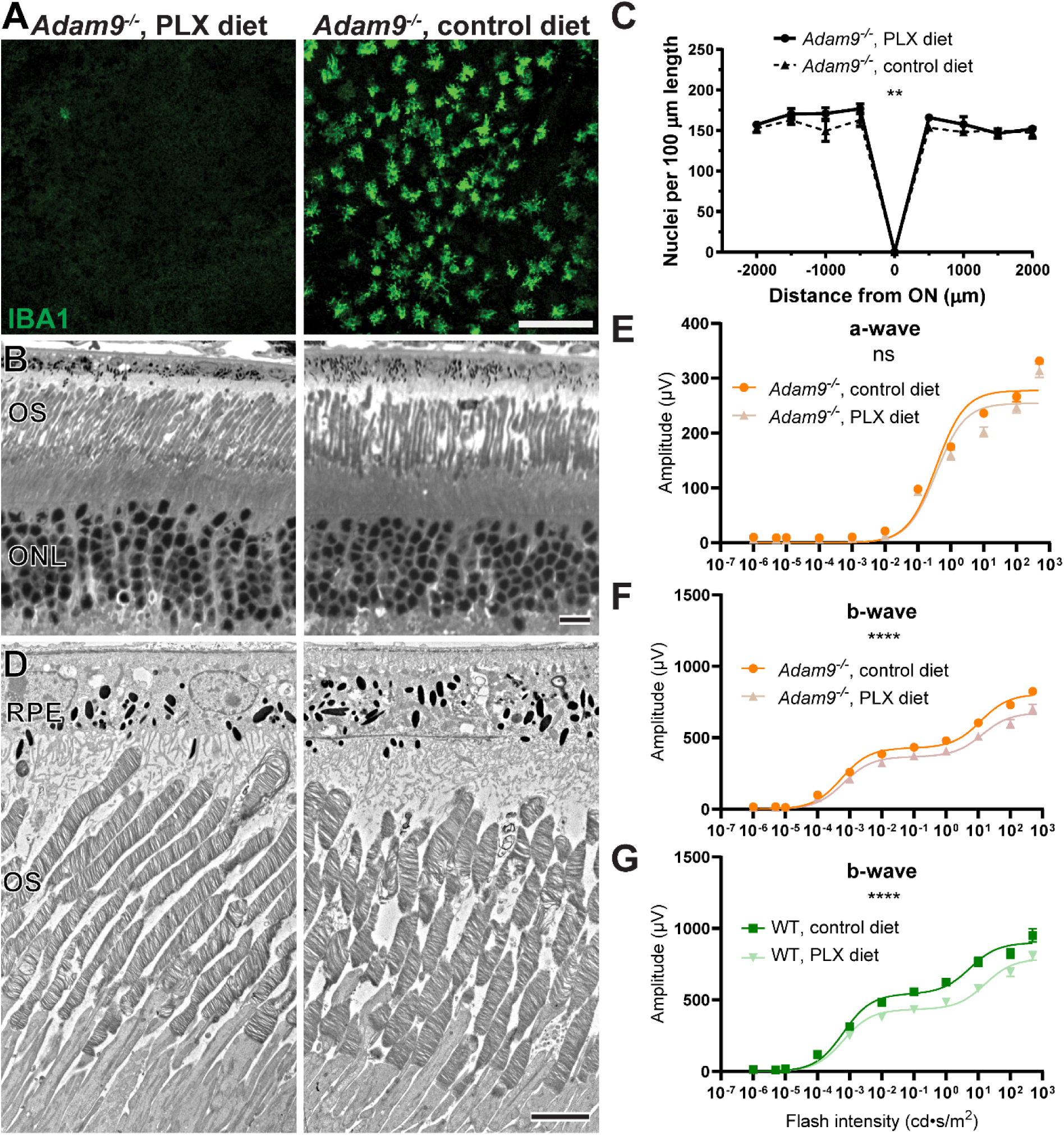
Ablation of subretinal mononuclear phagocytes does not affect the pathology associated with loss of ADAM9. **(A)** Representative immunofluorescence images of RPE flatmounts stained with IBA1 to label subretinal mononuclear phagocytes from P90 *Adam9^−/−^* mice fed a diet either with or without PLX3397 (PLX) beginning at P60. Scale bar: 100 µm. **(B)** Representative light microscopy images of *Adam9^−/−^* retinas treated with either control or PLX diet from P30 through P180. OS: outer segment; ONL: outer nuclear layer. Scale bar: 10 µm. **(C)** Quantification of the number of photoreceptor nuclei in 100 µm segments of retinal sections located at 500 µm intervals from the optic nerve (ON). For each condition, four mice were analyzed. Two-way ANOVA was used to determine that there was a statistically significant increase in photoreceptor numbers in PLX treated retinas compared to control retinas (p = 0.0090). **(D)** Representative TEM images of *Adam9^−/−^* retinas treated with either control or PLX diet. Scale bar: 5 µm. **(E)** ERG scotopic a-wave response of dark-adapted *Adam9^−/−^* mice treated with either control or PLX diet. For control diet, 18 eyes were analyzed. For PLX diet, 14 eyes were analyzed. Two-way ANOVA showed no statistically significant difference in genotype/light intensity (p = 0.1049). **(F)** ERG scotopic b-wave response of dark-adapted *Adam9^−/−^* mice treated with either control or PLX diet. For control diet, 18 eyes were analyzed. For PLX diet, 14 eyes were analyzed. Two-way ANOVA was used to determine that there was a statistically significant difference in genotype/light intensity (p < 0.0001). **(G)** ERG scotopic b-wave response of dark-adapted WT mice treated with either control or PLX diet. For control diet, 12 eyes were analyzed. For PLX diet, 16 eyes were analyzed. Two-way ANOVA showed a statistically significant difference in genotype/light intensity (p < 0.0001). Error bars represent mean ± SEM.

We next sought to analyze the effects of mononuclear phagocyte depletion on the pathology occurring in *Adam9^−/−^* mice. We treated these mice with PLX3397, beginning at P30, when IBA1-positive cells are first detected in the subretinal space, until P180, when significant photoreceptor degeneration has already occurred. We found that the ablation of these cells led to a very small yet statistically significant increase in the number of remaining photoreceptor nuclei (Figure 9, B and C), although there was no discernible change in the ultrastructure of photoreceptor cells or the subretinal space (Figure 9D). To determine whether PLX3397 treatment promoted the function of remaining photoreceptor cells, we performed ERG recordings from treated and untreated *Adam9^−/−^* mice and found no statistically significant difference in the amplitudes of a-waves produced by photoreceptors (Figure 9E). This may suggest that the small increase in the nuclear count in Figure 9C merely reflects the persistence of dead photoreceptor nuclei that are not cleared away by the mononuclear phagocytes. On the other hand, PLX3397 caused a small reduction in the amplitude of b-waves produced by bipolar cells downstream from photoreceptors (Figure 9F). Given that microglia play a homeostatic role in maintaining photoreceptor-to-bipolar cell synapses (24), we reasoned that this small decrease did not reflect the pathology taking place at the photoreceptor-RPE interface but rather may be attributed to the impaired maintenance of photoreceptor synapses in the absence of microglia. Consistent with this idea, we found that b-wave amplitudes of WT mice treated with PLX3397 were reduced to a similar degree as those in treated *Adam9^−/−^* mice (Figure 9G). Overall, we did not find any evidence to suggest that mononuclear phagocytes infiltrating the subretinal space of *Adam9^−/−^*mice significantly modulate the ongoing pathology.

## Discussion

The central finding of this study is that normal outer segment renewal can occur despite pathological expansion of the subretinal space, as evidenced by our findings with *Adam9^−/−^* mice. To sustain normal outer segment renewal under this condition, RPE cells form long pseudopods that extend across the expanded subretinal space to reach and ingest outer segment tips. This mechanism allows *Adam9^−/−^* photoreceptors to function normally and survive for long periods of time despite an early disruption of the subretinal space. One intriguing possibility is that the formation of elongated pseudopods represents an enhancement of the normal process of ingesting the tips of cone outer segments, which in WT mice are located at about half the height of rod outer segments. Once the subretinal space becomes expanded due to the knockout of ADAM9 or other related pathologies, elongated RPE pseudopods serve both cones and rods.

Despite the ability of the RPE to support outer segment renewal in *Adam9^−/−^* mice, photoreceptor cells in these mice eventually degenerate. In fact, the onset of photoreceptor degeneration occurs alongside a reduction in the number of pseudopods and a slowing of outer segment renewal. While it is conceivable that this degeneration is due to the continued expansion of the subretinal space to a point at which outer segment renewal becomes impaired, other pathological consequences of subretinal space expansion, such as an abnormal exchange of metabolites or retinoids, cannot be excluded as either primary or contributing factors. For example, altered photoreceptor metabolism was reported for another model characterized by an expanded subretinal space, the IMPG2 knockout mouse (25). Whether or not RPE pseudopods may facilitate transport of metabolites or retinoids, or the extent to which Müller glial cells can support both rod and cone photoreceptors following subretinal space expansion, remains a subject of future investigation.

Another finding of this study is that the knockout of ADAM9 is associated with a migration of activated mononuclear phagocytes to the subretinal space, a phenotype commonly observed in many models of photoreceptor degeneration (reviewed in (20)). Unique to the current study is that the expanded subretinal space of *Adam9^−/−^* mice allowed us to determine that these phagocytic cells were closely interdigitated with outer segment tips rather than with the RPE (e.g. Figure 8A). Furthermore, these cells contained outer segment material, indicating that they, in principle, have the ability to support outer segment turnover through the ingestion of outer segment tips. Yet, their pharmacological depletion led to no change in the ongoing pathology associated with loss of ADAM9, suggesting that the functional significance of any such outer segment ingestion may be of limited consequence.

There are several human conditions, such as vitelliform macular dystrophy and age-related macular degeneration associated with subretinal drusenoid deposits, in which patients exhibit an expansion of the subretinal space with visual function remaining reasonably preserved for long periods of time (7). In the case of vitelliform macular dystrophy, photoreceptors and the RPE could be separated by hundreds of microns (e.g. Figure 1). One intriguing explanation for why visual function persists despite the size of the vitelliform lesion is that RPE pseudopods may extend much further than observed in the *Adam9^−/−^* mouse. Yet testing this hypothesis awaits access to well-preserved human tissue from an early stage of disease.

Finally, we should discuss the limitations of this study, as well as further experimental directions that it opens. Although our findings in the *Adam9^−/−^* mouse primarily relate to the turnover of rod photoreceptors, vitelliform macular dystrophy is a disease that primarily affects cones (e.g. in Figure 1). The extent to which cone outer segment renewal is impaired by subretinal space expansion remains to be addressed in future experiments. Another interesting point is that, while the most prominent phenotype of the *Adam9^−/−^* mouse is pathological subretinal space expansion, *ADAM9* mutations in human patients lead to cone-rod dystrophy rather than vitelliform macular dystrophy. Assessing the role of ADAM9 in regulating the subretinal space and the photoreceptor-RPE interface in cone-dominant species would be invaluable in addressing this discrepancy. It would also be important to learn whether pseudopod formation occurs in other disease models of subretinal space expansion and how closely the extent of this expansion correlates with the rate of disease progression.

## Methods

### Sex as a biological variable

Mice of randomized sex were used in this study and sex was not considered as a biological variable.

### Statistics

For quantification of photoreceptor nuclei, two-way ANOVA was used. For ERG analysis, two-way ANOVA was used. For outer segment renewal analysis, unpaired t-test was used. For quantification of phagosomes and pseudopods, unpaired t-test was used. For quantification of rhodopsin, unpaired t-test was used. For single cell suction electrode recordings, unpaired t-test was used. All details of statistical analyses are further described in the corresponding figure legend. For figures, ns (not significant) represents p ≥ 0.05; * represents p < 0.05; ** represents p < 0.01, *** represents p < 0.001, **** represents p < 0.0001.

### Study approval

All patient data were obtained in compliance with an approved Duke University Health System Institutional Review Board protocol (Pro00092003). Written informed consent was received prior to participation, including for the use of photographs. Record of informed consent has been retained. Animal maintenance and experiments were approved by the Institutional Animal Care and Use Committees at Duke University (A184-22-10) and the University of California Davis (23883).

### Evaluation of human patient

Best-corrected visual acuity was measured using the Early Treatment Diabetic Retinopathy Study (ETDRS) retro-illuminated cabinet (Precision Vision). Macular spectral domain optical coherence tomography scans and fundus autofluorescence photographs were obtained using the Heidelberg Spectralis imaging platform. Microperimetry assessment was performed using the Nidek MP-1 microperimeter with a Goldmann III 200 ms stimulus. Multifocal electroretinogram recordings were obtained with the Diagnosys multifocal ERG with binocular LCD monitor using a 61-hexagon test and DTL electrodes. Genetic testing was performed by the Blueprint Genetics laboratory using next-generation sequencing.

### Animal husbandry

*Adam9^−/−^* mice were previously characterized in (14). WT mice (*Mus musculus*) were C57BL/6J (Jackson Labs stock #000664). All experiments were performed with animals of randomized sex. For drug treatment experiments, PLX3397 (pexidartinib) was purchased from Medkoo (206178) and compounded into the Open Standard Diet by Research Diets (D11112201i) at a concentration of 400 mg PLX3397 per kg of chow. The Open Standard Diet served as the control diet.

### Plastic block embedding for light microscopy, TEM and 3D-ET

At 1.5 hours after light onset, anesthetized mice were transcardially perfused with a solution of 2% paraformaldehyde, 2% glutaraldehyde and 0.05% calcium chloride in 50 mM MOPS (pH 7.4). Enucleated eyes were fixed for an additional 2 hours in the same fixation solution at RT. Eyes were dissected and cut in half through the optic nerve. Tissue was treated with 2% osmium tetroxide (Electron Microscopy Sciences), dehydrated with ethanol and infiltrated and embedded in Embed 812 resin (Electron Microscopy Sciences).

### Light microscopy of plastic sections

500 nm thick retinal sections were cut and stained with methylene blue. Images were taken with a confocal microscope (Eclipse 90i and A1 confocal scanner; Nikon) with a 60× objective (1.49 NA Plan Apochromat VC; Nikon) and NIS-Elements software (Nikon). Image analysis and processing was performed with ImageJ. For quantifying photoreceptor nuclei, nuclei were counted in 100 µm boxes at 500 µm intervals from the optic nerve spanning 2,000 µm in each direction.

### Transmission electron microscopy (TEM)

70 nm sections were cut, placed on copper grids and counterstained with 2% uranyl acetate and 3.5% lead citrate (19314; Ted Pella). The samples were imaged on a JEM-1400 electron microscope (JEOL) at 60 kV with a digital camera (BioSprint; AMT). Image analysis and processing was performed with ImageJ.

### 3-Dimensional electron tomography (3D-ET)

Serial 500 nm thick retinal sections were cut and placed on 50 nm Luxel film slot grids. Grids were glow-discharged on both sides, and a mixture of 10 nm, 20 nm and 60 nm gold particles were deposited on the sample surfaces to serve as fiducial markers. 3D-ET was conducted on a Titan Halo (FEI) operating at 300 kV in STEM mode. A four-tilt series data acquisition scheme previously described (26) was followed in which the specimen was tilted from −60° to +60° every 0.5° at four evenly distributed azimuthal angle positions. Images were collected with a high-angle annular dark field (HAADF) detector in STEM mode. The final volumes were generated using an iterative reconstruction procedure (26). 3dmod and ImageJ were used for image analysis. Segmentation was manually obtained using histogram-based utilities in the Amira-Avizo software suite.

### Electroretinography (ERG)

ERGs were recorded using the Espion E2 system with a ColorDome Ganzfeld stimulator (Diagnosys LLC). Following dark-adaptation, mice were anesthetized by an intraperitoneal injection of 100 mg/kg ketamine and 10 mg/kg xylazine. Pupils were dilated with a mixture of 1% cyclopentolate HCl and 2.5% phenylephrine. Eyes were lubricated during the recordings with 2.5% hypromellose and body temperature was maintained by a heated platform. Recordings were made using gold contact lens electrodes (Mayo Corp) with stainless steel needle electrodes (Ocuscience) in the mouth (reference) and at the base of the tail (ground). ERG signals were sampled at 1 kHz and recorded with 0.15-Hz low-frequency and 500-Hz high-frequency cutoffs. Responses to flashes from 0.00001 to 500 cd·s/m^2^ with 1–10 trials (fewer trials for the brighter stimuli) averaged and interflash intervals of 5–180 s were recorded in the dark. The data from the ERG recordings were analyzed using MATLAB 2016a (MathWorks) as described in (27, 28). Oscillatory potentials were removed from the signals by 55-Hz fast Fourier transform low-pass frequency filtering. The amplitude of the b-wave was calculated from baseline to the peak for dimmer flashes and from the bottom of the a-wave to the b-wave peak for brighter flashes. Data points from the a-wave and b-wave stimulus–response curves were fitted by single- or double-hyperbolic functions respectively, using the least-square fitting procedure.

### Intravitreal injection of CF-568-hydrazide

Mice at P30 and P180 were anesthetized with 1.5% isoflurane and topical application of 0.5% proparacaine HCl solution. Eyes were injected intravitreally with 2 µl of 0.5% CF-568-hydrazide in PBS. Following the injection, topical erythromycin ointment was applied to the ocular surface, and mice were returned to normal housing after recovery from anesthesia. At 6 days post-injection, mice were euthanized and retinas were harvested in mouse Ringer’s solution (containing 130 mM NaCl, 3.6 mM KCl, 2.4 mM MgCl_2_, 1.2 mM CaCl_2_, and 10 mM HEPES, pH 7.4). Outer segments were mechanically dissociated, fixed in 4% paraformaldehyde in PBS, and subsequently stained with fluorescent-conjugated wheat germ agglutinin (WGA) (1 µg/mL; W11261; Thermo Fisher Scientific). Outer segments were imaged on coverslip bottom dishes (MatTek) using the Nikon confocal microscope described above. Image analysis and processing was performed with ImageJ. The length from the outer segment base (indicated by either a small remaining portion of the inner segment or WGA staining of the newly forming discs or connecting cilium) to the CF-568-hydrazide band was measured and divided by the total length of the isolated outer segment to generate the fraction of outer segment length that the CF-568-hydrazide band travelled over 6 days.

### Rhodopsin difference spectroscopy

Mice at P30 were dark-adapted overnight before removing the cornea and lens under dim red light. Eyecups were lysed in 600□µl of 2% octyl glucoside with protease inhibitor (cOmplete; Sigma Aldrich) and sonicated 3 times each of 5□seconds. Lysates were centrifuged at 10,000 × g for 10□minutes and the rhodopsin concentration of the supernatant was measured by difference spectroscopy (29). 25 mM hydroxylamine, pH 7.5 was added to 100□μl of lysate and the initial spectra was recorded from 300 to 700□nm using a DU800 spectrophotometer (Beckman). The sample was then bleached with a 100-W halogen lamp for 60□s before taking the second spectrum reading. The difference in the absorbance at 500□nm was used to calculate rhodopsin concentration, using the extinction coefficient of 40,500□M^−1^ ·cm^−1^.

### Single cell suction electrode recordings

Suction electrode recordings from the outer segments of intact mouse rods were performed as previously described (30). Briefly, P30 mice were dark-adapted overnight, euthanized, and their retinas were dissected and stored on ice in L-15 medium supplemented with 10□mM glucose. Recordings were performed in oxygenated, bicarbonate buffered Locke’s solution supplemented with 10 mM glucose at 35–37□°C. A suction pipette containing HEPES-buffered Locke’s solution (pH 7.4) recorded the electrical responses to brief (10□ms, 500□nm) flashes of calibrated strength, which were amplified (Axopatch 200B; Molecular Devices), filtered at 30 Hz with an eight-pole Bessel (Frequency Devices), and digitized at 200□Hz using custom-written acquisition procedures in IgorPro (Wavemetrics). Responses to a large number (>30) of dim flashes were averaged and used to determine the mean time to peak, exponential time constant of the recovery phase (*τ*_rec_)and integration time (time integral of the response divided by the peak amplitude). Saturating responses to bright flashes were used to calculate the dominant time constant of recovery (τ_D_), as previously described (30–32). Flash sensitivity (I_o_) was defined as the flash strength that suppressed one-half of the circulating dark current (I_dark_).

### Immunofluorescence microscopy of RPE flatmounts

Anesthetized mice were transcardially perfused with a solution of 4% paraformaldehyde in PBS. Prior to enucleation, a cautery pen was used to create a small superficial burn on the superior cornea. Enucleated eyes were fixed for an additional 2 hours in the same fixation solution at RT. The cornea was removed apart from a small flap on the superior region to identify orientation. The lens was removed and four cuts were made in the eyecup to produce superior, inferior, nasal, and temporal regions. The retina was then removed and the remaining eyecup consisting of sclera and RPE was processed for immunofluorescence staining. Tissue was blocked in PBS containing 7% donkey serum and 0.5% Triton X-100 for 30 min at RT and incubated with primary antibodies, including rabbit anti-IBA1 (1:1000; 019-19741; Wako) and/or rat anti-CD68 (1:500; 137002; BioLegend), overnight at 4°C. Tissue was washed and incubated with secondary donkey anti-rabbit and anti-rat antibodies conjugated to Alexa Fluor 488 or 647 (1:1000; A21206 and A78947, respectively; Thermo Fisher Scientific), Alexa Fluor 568 phalloidin (1:400; A12380; Thermo Fisher Scientific) and Hoechst 33342 (10 µg/mL; 62249; Thermo Fisher Scientific) for 2 hr at RT. Tissues were washed and mounted onto slides with Shandon Immu-Mount (9990402; Thermo Fisher Scientific). Images were acquired using the Nikon confocal microscope described above with a 10× objective (0.45 NA Plan Apochromat λ; Nikon). The large image tool was used to capture a 6 x 6 field that encompassed the entire eye in an appropriate z depth for each sample (estimated range about 70-100 µm depending on the sample) at a 12 µm step size and a pixel resolution of 1024 × 1024. Single z images were captured with a 20× objective (0.75 NA Plan Apochromat λ; Nikon). Images were processed in ImageJ and shown as maximum intensity z projections.

### Data availability

Values for all data points in graphs are reported in the Supporting Data Values file.

## Author contributions

TRL contributions included: designing research studies, conducting experiments, analyzing data and writing and editing the manuscript.

CMC contributions included: conducting experiments, analyzing data and editing the manuscript.

SP contributions included: conducting experiments, analyzing data and editing the manuscript.

CRS contributions included: conducting experiments, analyzing data and editing the manuscript.

KKH contributions included: conducting experiments, analyzing data and editing the manuscript.

K-YK contributions included: conducting experiments, analyzing data and editing the manuscript.

MHE contributions included: designing research studies, analyzing data and editing the manuscript.

OA contributions included: clinical analysis, conducting experiments, analyzing data and editing the manuscript.

MEB contributions included: designing research studies, analyzing data and editing the manuscript.

VYA contributions included: designing research studies, analyzing data and writing and editing the manuscript.

## Acknowledgements

This work was supported by the National Institutes of Health grants EY033763 (TRL), EY030451 (VYA), EY005722 (VYA), EY033857 (OA), EY024320 (MEB), U24NS120055 (MHE), S10OD034447 (MHE), Duke University Physician-Scientist Strong Start Award (OA), Research to Prevent Blindness Career Development Award (OA) and an Unrestricted Award from Research to Prevent Blindness Inc. (Duke University). This work is subject to the NIH Public Access Policy. Through acceptance of this federal funding, the NIH has been given a right to make the work publicly available in PubMed Central.

## Supplemental material

**Supplemental Figure 1.**
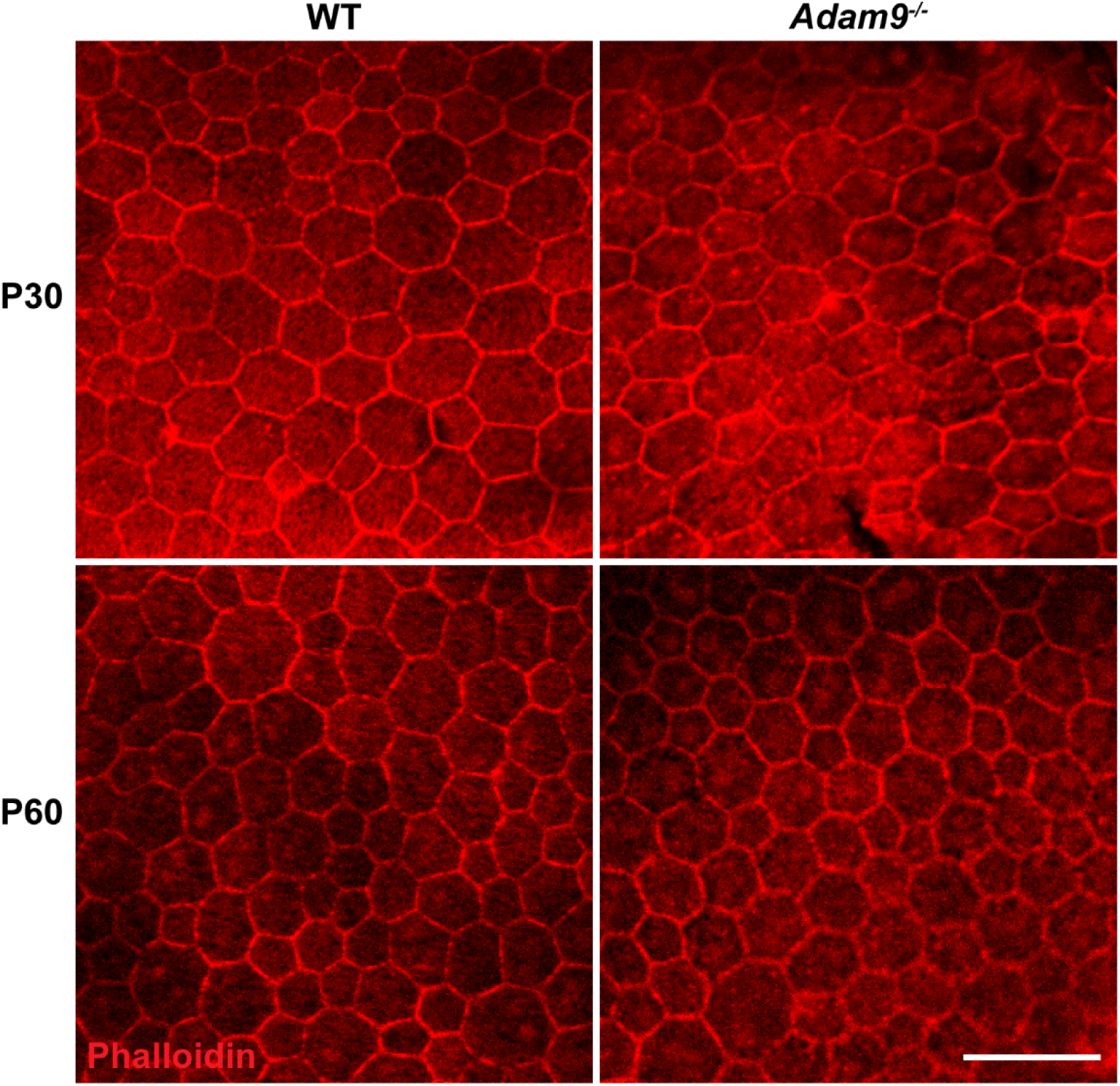
Loss of ADAM9 does not lead to overt pathology in the RPE monolayer at early timepoints. Representative immunofluorescence images of RPE flatmounts stained with phalloidin to visualize RPE cell borders from WT and *Adam9^−/−^* mice at P30 and P60. Scale bar: 50 µm.

**Supplemental Figure 2.**
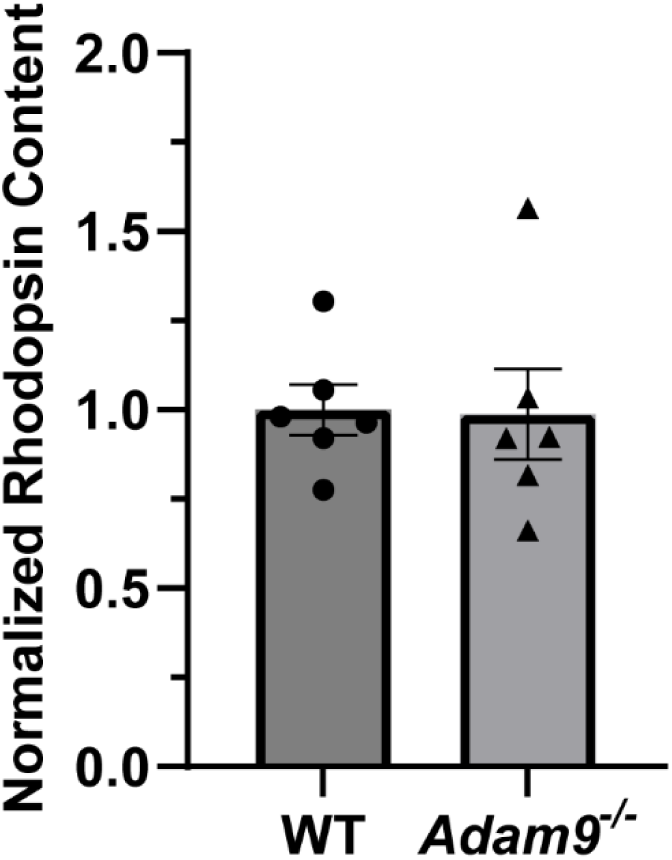
Loss of ADAM9 does not affect the total rhodopsin content in the retina prior to photoreceptor degeneration. Total rhodopsin content of dissected eyecups was determined by difference spectroscopy and normalized to the average WT value. Unpaired t-test showed no statistically significant difference in rhodopsin content between WT and *Adam9^−/−^* eyecups at P30 (p□=□0.9321). For each genotype, six eyecups were analyzed. Error bars represent mean ± SEM.

**Supplemental Figure 3.**
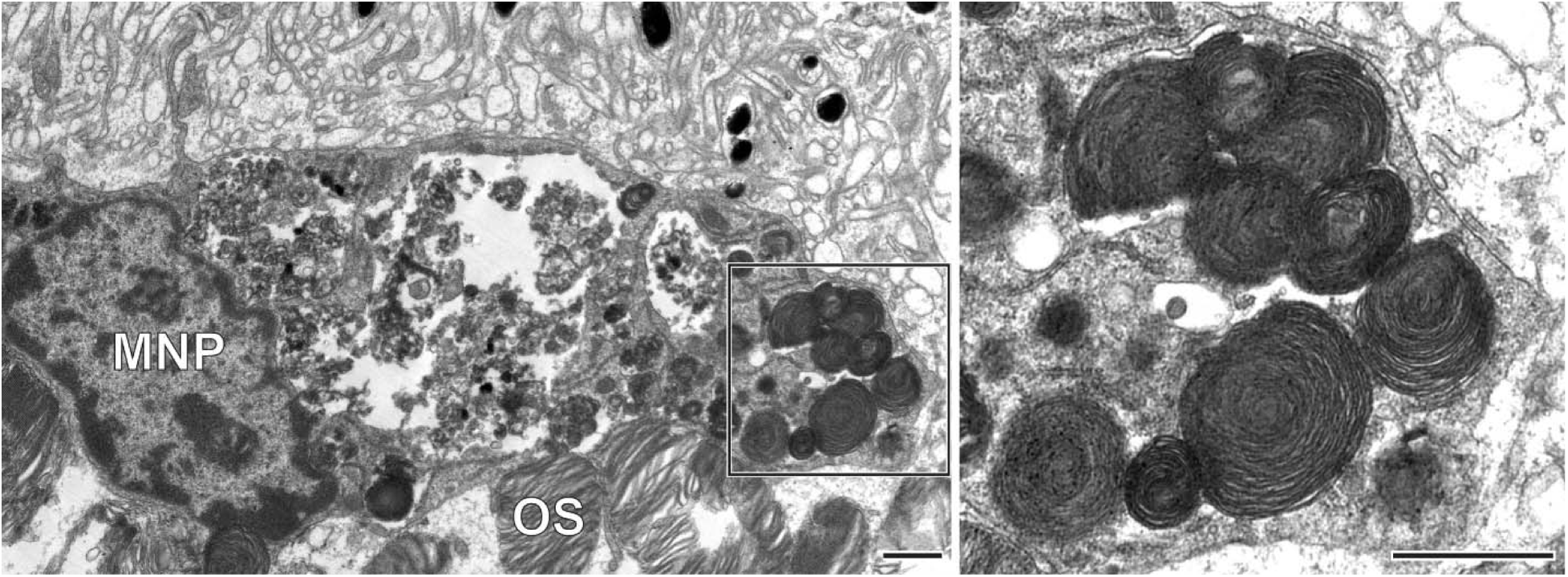
Subretinal mononuclear phagocytes are capable of ingesting outer segment material. Representative TEM images of a mononuclear phagocyte (MNP) in the subretinal space of an *Adam9^−/−^* mouse at P180. Boxed area containing several phagosomes is shown at higher magnification to the right. OS: outer segment. Scale bars: 1 µm.

**Movie 1. Reconstructed tomograms across four serial sections from an *Adam9^−/−^* retina – Example 1**

Shown is an ∼2 µm thick volume of the *Adam9^−/−^* retina with an isotropic resolution of 7.4 nm.

**Movie 2. Reconstructed tomograms across four serial sections from an *Adam9^−/−^* retina – Example 2**

Shown is an ∼2 µm thick volume of the *Adam9^−/−^* retina with an isotropic resolution of 7.4 nm.

